# Droplet-based Random Barcode Transposon-site Sequencing (Droplet RB-TnSeq) to characterize phage-host interactions

**DOI:** 10.1101/2025.09.29.679331

**Authors:** Fangchao Song, Allison Hung, Madeline Moore, Vivek Mutalik, Adam P. Arkin

**Affiliations:** Environmental Genomics & Systems Biology Division, Lawrence Berkeley National Laboratory, Berkeley, CA, US; Department of Molecular and Cell Biology, University of California at Berkeley, Berkeley, CA, US; Department of Bioengineering, University of California at Berkeley, Berkeley, CA, US

**Keywords:** droplet RB-TnSeq, bacteriophage T4 and N4, phage resistance, droplet microfluidics, random barcode transposon-site sequencing (RB-TnSeq)

## Abstract

Bacteriophages (phages), the most abundant self-replicating entities on Earth, are central to microbial ecology and hold promise as therapies against antibiotic-resistant pathogens. However, the molecular determinants of phage adaptation to hosts remain poorly defined. While bulk genetic screens such as transposon sequencing are high throughput, running all mutants in mixed populations biases outcomes toward phage receptor discovery. Competition among resistant mutants and the escalating multiplicity of infection (MOI) from additional phages released from lysed cells often mask moderately acting host factors—such as inner-membrane and energy-transduction systems supporting DNA entry, surface modifiers, and regulators of host defense. To overcome these limitations, we developed droplet-based random barcode transposon-site sequencing (Droplet RB-TnSeq), which encapsulates single barcoded mutant cells with defined phage titers. Applying this method to *Escherichia coli* infected with phages T4 and N4, we reproducibly recovered known determinants (*ompC* for T4 and *nfrA/B* for N4) and identified additional contributors, including membrane proteins, polysaccharide modifiers, signaling modules, and several uncharacterized genes. Notably, mutations in *chbR/C/F*, *ygaZ/H*, and *yfbO/L* were found to confer resistance to N4, validated by plaque assays and complementation. Droplet RB-TnSeq thus enables genome-wide dissection of host factors beyond receptor-level interactions under well-controlled infection conditions.

## Introduction

Bacteriophages (phages) - the viruses of bacteria, are the most abundant organisms in the biosphere.^1^ They play essential roles in the composition and function of microbial communities, thereby influencing human and plant health, and climate change.^2–4^ Due to the high efficiency and specificity to their host bacteria, phage therapy is considered the most promising alternative to antibiotics in the age of multidrug resistance.^5,6^ It is vital to understand the mechanism of phage infection and bacterial responses to diverse phages for understanding their ecological impact and designing phage therapy to combat the antibiotic resistant crisis. However, although current advances in whole genome sequencing and the availability of bacterial mutants and phage collections have expanded our understanding of phage-host interactions,^7,8^ many mechanisms of phage infection and bacterial host factors are still unknown due to the complexity of phage-host interactions.^9^ During the phage infection, phage may rely on or be mediated by various host factors including phage receptors, metabolic factors, DNA translocation co-factors, surface polysaccharide/biogenesis modifiers, and multiple diverse host defense mechanisms.^10,11^ It is imperative to have a scaled-up comprehensive genetic screening to rapidly discover new mechanisms of phage-host interaction and phage resistance.

Techniques like RB-TnSeq, Dub-Seq, and CRISPRi increase the ability to approach new host-phage pairs and enable discovery of host factors that impact phage infection at genome scale by combining loss-of-function or overexpression libraries with high throughput sequencing.^7,12–15^ However, these bulk competitive assays suffer from bias. First, all the mutants compete for the shared nutrients for growing, the highest resistant host mutants would outcompete others and inhibit the growth of other milder resistant mutants.^16,17^ Second, the susceptible mutants will release more phages into the environments, increasing the selective pressure.^18^ It will further suppress the milder resistant strains. This results in detection of the highest resistant host mutants, such as phage’s host receptors, but masks more nuanced determinants. Droplet microfluidics is a potential solution for solving this issue by removing competition through isolation of bacteria-phage pairs. Using customized microfluidic chips and well controlled cell density, millions of water-in-oil droplets with single microbial cells can be generated with defined environments such as phages and antibiotics.^19,20^ Each droplet serves as a reactor that allows individual strains of microbial cells grow and prevents the competition amongst different variants.^21^ Due to the unique high throughput ability of single cell isolation and partition, droplet microfluidics has been successfully used for various single cell analysis, such as RNAseq and ATAC-seq.^22–24^ Notably, Thibault et al.^25^ demonstrated that combining single-cell droplet technology with traditional transposon mutant library techniques—referred to as droplet Tn-Seq—enables isolated cell growth, microscopy, and FACS, revealing distinct growth profiles between bulk Tn-Seq and droplet Tn-Seq.

Here, we develop droplet-based random barcode transposon-site sequencing (Droplet RB-TnSeq) to assess gene functions of microbes during phage infection by encapsulating single cell from the RB-TnSeq microbial library with a controlled titer of phage. Each mutant grows in its own droplet microreactor without interference from others. This method extends the Droplet Tn-Seq, but has much simpler sample preparation and gene fitness measurement compared to Droplet Tn-Seq. We demonstrate that Droplet RB-TnSeq reproducibly and reliably uncovers the factors influencing microbial susceptibility to phages. Not only identifying all the known factors, Droplet RB-TnSeq also finds new genes that affect phage susceptibility. Those genes may reveal novel mechanisms of phage infection and host factors.

## Results

### Droplet RB-TnSeq

We used *E. coli* Keio ML9a strain and phages N4 and T4 as model systems to benchmark the assay platform. Both phage N4 and T4 are highly distinct, employing different mechanisms for host recognition, replication, and cell lysis efficiency. In the Droplet RB-TnSeq assay (*Figure 1A*), an *E. coli* Keio ML9a RB-TnSeq library^15^ in LB medium and *E. coli* phage solution were mixed in a 2-inlet microfluidic chip and encapsulated into water-in-oil droplets (*Figure S1*). By controlling the cell densities of input *E. coli* and phage solutions as well as the size of droplets, single cells (for occupied droplets) were encapsulated in individual droplets with growth medium and phages. The droplet size was controlled at 60-um diameter in this experiment, which provides sufficient space to hold substantial bacterial cells (>900 cells per droplet) after growth while still maintaining thermodynamic stability during incubation. The phage number per droplet was determined by Poisson distribution driven by random and constant probability of droplet compartmentation. Phage titer of the input phage solution was assessed by phage plaque assay test one day before use.

**Figure 1.**
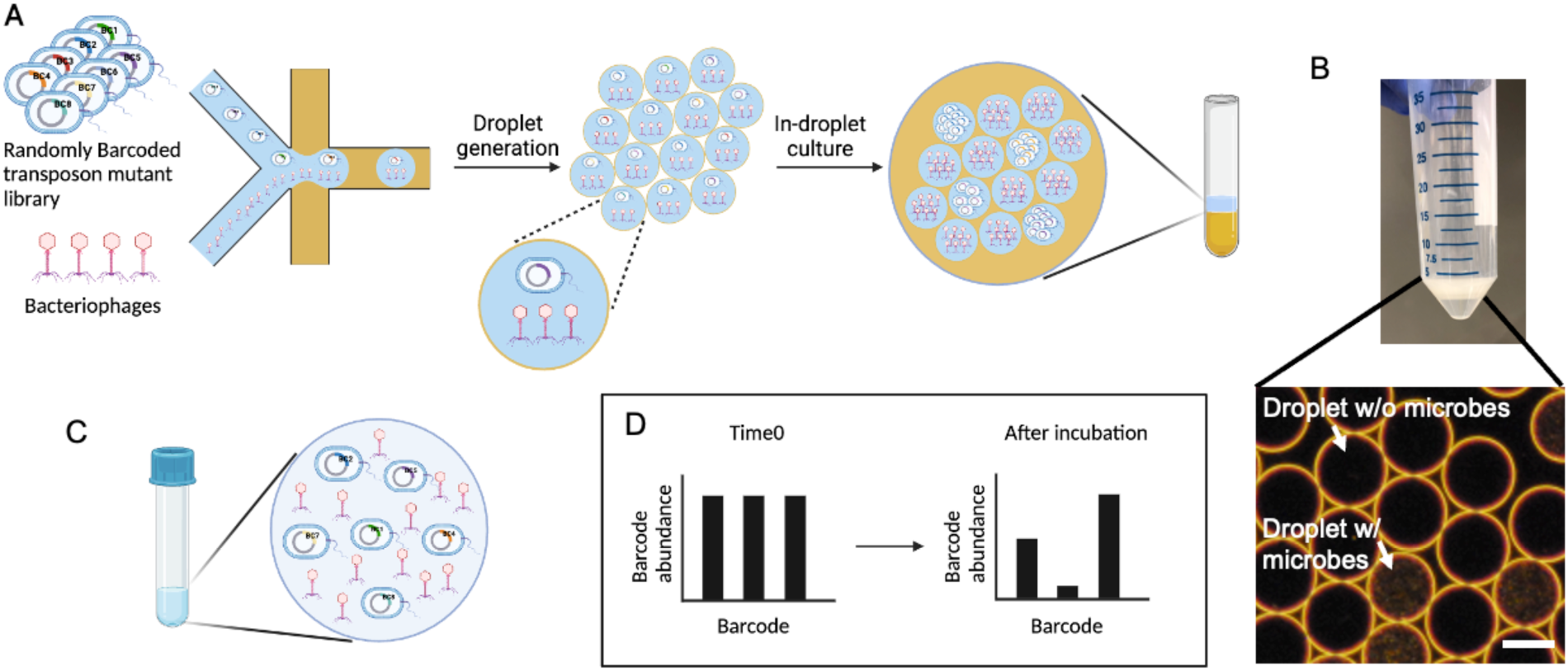
Overview of Droplet RB-Tnseq and bulk RB-TnSeq. (A) Droplet RB-TnSeq. Randomly barcoded loss-of-functions transposon mutant library strain and bacteriophages are mixed and encapsulated into water-in-oil droplets. Single host mutant strain is incubated with phages. And the t0 and t samples during incubation are sequenced. (B) The representative image of droplet cultures. (Bar = 50 um) (C) Bulk RB-TnSeq. Hundreds of thousands of randomly barcoded loss-of-function transposon mutant library strains are mixed with bacteriophages and incubated in one tube. And the t0 and t samples during incubation are sequenced. (D) Fitness calculation by RB-TnSeq. The fitness of each strain is calculated based on the barcode abundance at the start point at t0 and after incubation at t.

Two input concentrations of phages were used in this study. One was 4.4 × 10^7^ pfu/mL for an overall MOI equal to 5 (refer to “LMOI” in this work); another was 5.1 × 10^9^ pfu/mL for an overall MOI equal to 580 (refer to “HMOI” in this work). The microfluidic chip was fabricated using PDMS with a mode manufactured using SU-8 photolithography. The design is shown in Figure S1 (dwg file is in the supplementary data). After the droplets were generated, the bacteria and phages in droplets were incubated in a culture tube (*Figure 1B*). If the mutant was resistant to the phage, it would grow in the droplets, thereby increasing fitness. Otherwise, the single bacterial cell would die in the droplets. The droplets were intact after incubation as shown in *Figure 1B*. As a comparison, *Figure 1C* represents the Bulk RB-TnSeq assay, in which all the bacterial cells and phages were cultured in one mixture in a tube. The abundances of each mutant in both Droplet RB-TnSeq and Bulk RB-TnSeq were determined by Barseq. Fitness scores were calculated by comparing the abundance changes of each individual mutant to the average abundance change of all the mutants (*Figure 1D*), following the same protocol previously described.^15^ Higher fitness during phage infection indicates more resistance to phages.

### Validation and optimization of Droplet RB-TnSeq

The Droplet RB-TnSeq assay starts with the encapsulation of single RB-TnSeq microbial library cells into droplets and ends with Barcode sequencing (Barseq) of the recovered microbial DNA from the water-in-oil emulsions. To ensure those steps are unbiased in representing the microbial composition, we first compared the relative abundance of the original input RB-TnSeq library (OD_600_ = 1) to the relative abundance of the droplet microbial single cell encapsulations without cultivation. Briefly, we generated single cell microbial droplet cultures from 1 mL *E. coli* RB-TnSeq library cultures with cell density of 8.8 × 10^5^ cell/mL, which was diluted from the original input culture of OD_600_ = 1. The collected droplet culture was immediately processed to remove the oil and extract the DNA without cultivation. The relative abundance of both original input culture (OD_600_ = 1) and uncultured droplet culture were measured by Barseq and shown in *Figure 2A*. The results agree with each other well (R^2^ = 0.99), indicating that the droplet generation and DNA extraction from emulsion would not bring in measurement bias.

**Figure 2.**
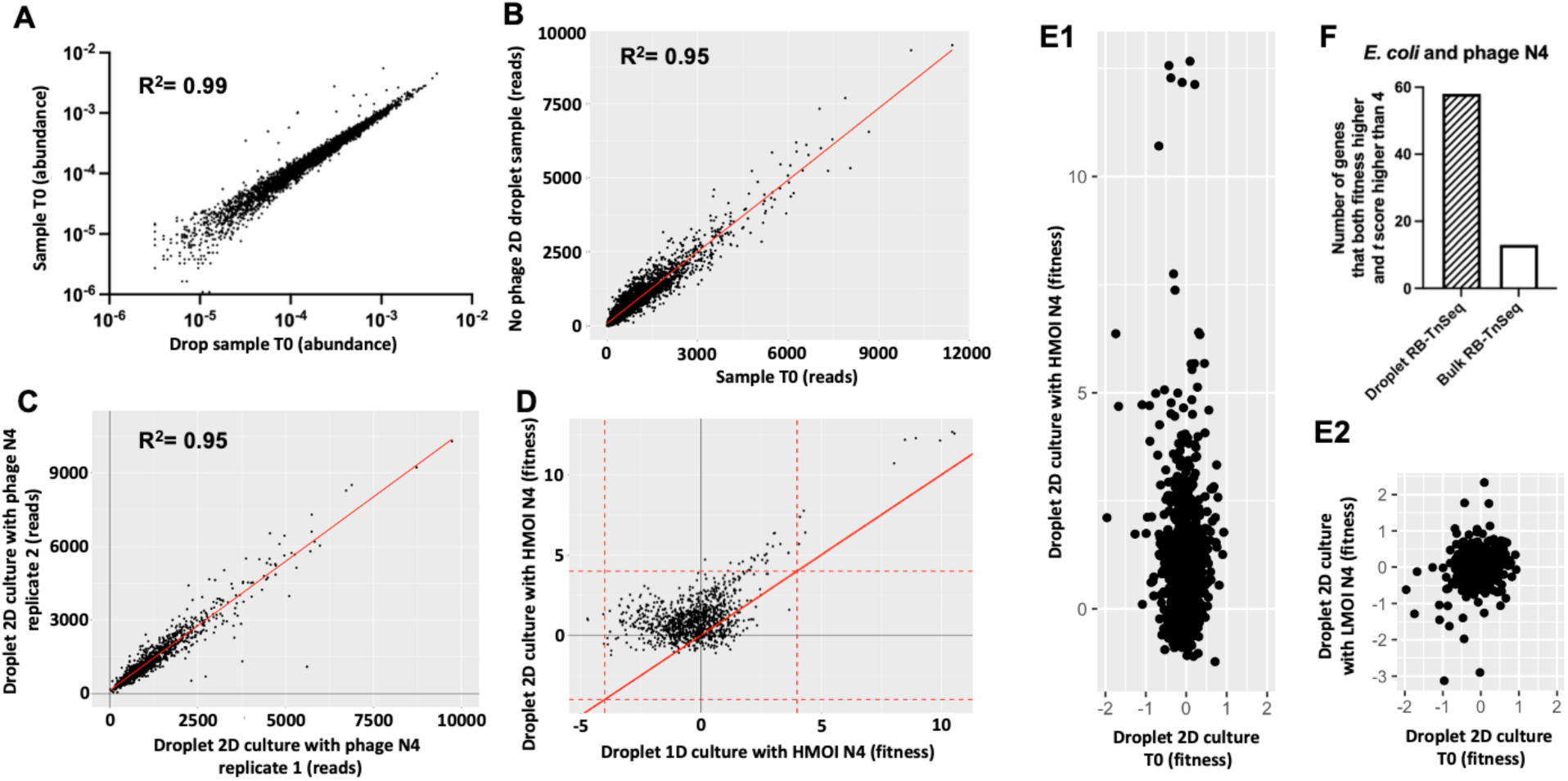
Validation of Drop RB-TnSeq. (A) The comparison of the abundance of *E. coli* mutants between the original bulk sample and the droplet sample without cultivation. (B) The comparison of the abundance of *E. coli* mutants between the original bulk sample and the droplet sample cultivated for 2 days without phages. (C) The abundance of two replicates of *E. coli* mutant droplet sample cultivated for 2 days with phage N4. (D) The comparison of the fitness scores of *E. coli* mutant droplet samples cultivated for 1 day and 2 days with phage N4. (E1) The comparison of the fitness scores of *E. coli* mutants between the droplet samples cultivated for 2 days with High MOI phage N4 and the droplet samples without cultivation. (F2) The comparison of the fitness scores of *E. coli* mutants between the droplet samples cultivated for 2 days with Low MOI phage N4 and the droplet samples without cultivation. (F) The number of *E. coli* mutants that has both fitness score and *t* score higher than 4 from Droplet RB-TnSeq and Bulk RB-TnSeq with phage N4.

The droplet reactors – 60-um diameter droplets surrounded by oil and biocompatible surfactant have higher surface to volume ratio compared to flasks or culture tubes. To investigate if the cell growth in this small, high surface-to-volume ratio droplet reactor would influence our measurements, we compared the relative abundance of the original input RB-TnSeq library (OD_600_ = 1) with the relative abundance of the droplet microbial single cell culture that is cultured for 2 days in LB without phages. All the cells are expected to grow to stationary phase in their own droplets, resulting in relative abundances that mirror the input culture, expect where specific growth defects affect particular mutants. *Figure 2B* represents this result. The number of reads of each mutant in the cultured droplet *E. coli* RB-TnSeq library is consistent with the number of reads of each mutant in the original input cultured droplet *E. coli* RB-TnSeq library (R^2^ = 0.95), indicating that the cells grow well in water-in-oil droplet reactor with LB and that the relative abundances do not change. To further investigate the reliability of Droplet RB-TnSeq, we conducted Droplet RB-TnSeq using the *E. coli* Keio ML9a RB-TnSeq library and phage N4 with two replicates. The number of reads of each mutant in each replicate is plotted in *Figure 2C*. The two replicates agree with each other (R^2^ = 0.95), indicating that Droplet RB-TnSeq is replicable.

Droplet RB-TnSeq can be optimized for studying microbial phage resistance through varying experimental parameters. One parameter is the cultivation time. Although most lytic phages are very effective on lysing host cells in hours, we incubate the co-cultures over days to target the slow growing survival mutants. 1-day (1D) and 2-day (2D) microbe-phage N4 co-cultures (HMOI) were compared. The fitness scores obtained from the Barseq analysis of both 1D and 2D cultures are plotted in *Figure 2D*. The red solid line indicates the same fitness in both 1D and 2D cultures. And the red dot lines indicate fitness scores equal to -4 and 4. The fitness scores of most mutants in 2D culture is higher than 1D culture. In particular, mutants with high fitness scores (>4 in either culture, indicating strong resistance upon loss of the cognate gene) all have higher fitness scores in 2D culture than 1D culture. It suggests that 2D culture that allows longer surviving host cell growth gives better resolution on identifying phage resistant mutants.

A second key parameter is the multiplicity of infection (MOI)—the ratio of phages to bacterial cells. In bulk culture, MOI is easily controlled by manually mixing defined concentrations of phages and bacteria. In contrast, during high-throughput droplet generation—producing up to 6 million samples per hour—achieving uniform MOI across all droplets is not feasible due to the stochastic nature of Poisson loading. To ensure single-cell resolution, we deliberately use a low bacterial density, resulting in ∼90% of droplets being empty, while the remaining droplets contain exactly one bacterial cell. However, the droplet generation process cannot guarantee uniform phage loading at this level, instead producing a distribution of phage counts across droplets [*Figure S2*]. Consequently, the droplets contained a single bacterial cell, and the phage loading followed a Poisson distribution centered around the average MOI. This variability can lead to inconsistent phage infection efficiencies—particularly at low MOIs (less than 4, *p* = 0.26), which may fail to lyse the bacterial cells, potentially resulting in false positives in the assay. To mitigate this issue, a high MOI (HMOI) droplet loading strategy was also employed. During droplet culture generation, the bacterial input density was kept the same as in the low MOI (LMOI) condition to achieve single bacterial cell loading, while the phage input density was increased to on average 580 phages per droplets, resulting in an average MOI of 580 with no droplets having less than 450 phages (*p* = 10^-8^) [*Figure S2(B)*]. The absence of droplets with low phage numbers minimizes the likelihood of false positives, which may arise under the LMOI condition. Therefore, resistance phenotypes measured under the HMOI sample are more likely to reflect true phage resistance. This is supported by the data in *Figure 2E*, which compares the fitness scores of each mutant after a 2D droplet culture under HMOI (*Figure 2E1*) or LMOI (*Figure 2E2*) with those from the initial droplet sample (T₀). The HMOI condition shows a broader range of fitness scores, with some mutants exhibiting strong resistance (fitness > 10). In contrast, fitness scores under the LMOI condition closely resemble those of the T₀ sample, with more uniform growth across mutants—likely due to the presence of droplets with insufficient phage numbers to challenge bacterial cells. Based on these findings, the HMOI condition is recommended for Droplet RB-TnSeq assays. All droplet-based phage resistance data presented in this study were obtained under HMOI samples.

To validate the advantage of Droplet RB-TnSeq assay compared to Bulk RB-TnSeq assay, we compare the number of mutants with both high fitness score that indicates the mutant growth and *t* score that indicates the confidence of the fitness score (both > 4) in Droplet RB-TnSeq and Bulk RB-TnSeq. *The E. coli* Keio ML9a RB-TnSeq mutant library was challenged by phage N4 using both Droplet RB-TnSeq and Bulk RB-TnSeq. Single *E. coli* mutant cells were cultivated with phage N4 in droplets at the HMOI condition; and the pooled *E. coli* Keio ML9a RB-TnSeq mutants were cultivated with phage N4 at high MOI (MOI 234). The results are shown in *Figure 2F*. Using the cutoff of fitness score higher than 4 and *t* score higher than 4, we revealed 58 mutants in Droplet RB-TnSeq, but only 13 mutants in Bulk RB-TnSeq, which is 4 times less than Droplet RB-TnSeq. The top hits from Droplet RB-TnSeq are shown in *Figure 5B*.

### Revealing the resistance of *E. coli* on phage T4 using Droplet RB-TnSeq

Given we have determined optimized conditions —high multiplicity of infection (HMOI) and a two-day incubation period (2D), we now apply them to investigate the impact of *E. coli* genes on phage T4 infection. *Figure 3A* shows the fitness and *t* scores of *E. coli* Keio ML9a RB-TnSeq mutants in the presence of phage T4. Genes with fitness > 4 and |*t* score| > 4 are highlighted in green and listed in *Figure 3B*. Aside from a few previously known phage T4-resistant genes (*ompC*, *ompR*, *envZ*) - marked as pink in *Figure 3B*, most of the identified genes have not been previously associated with T4 resistance.

**Figure 3.**
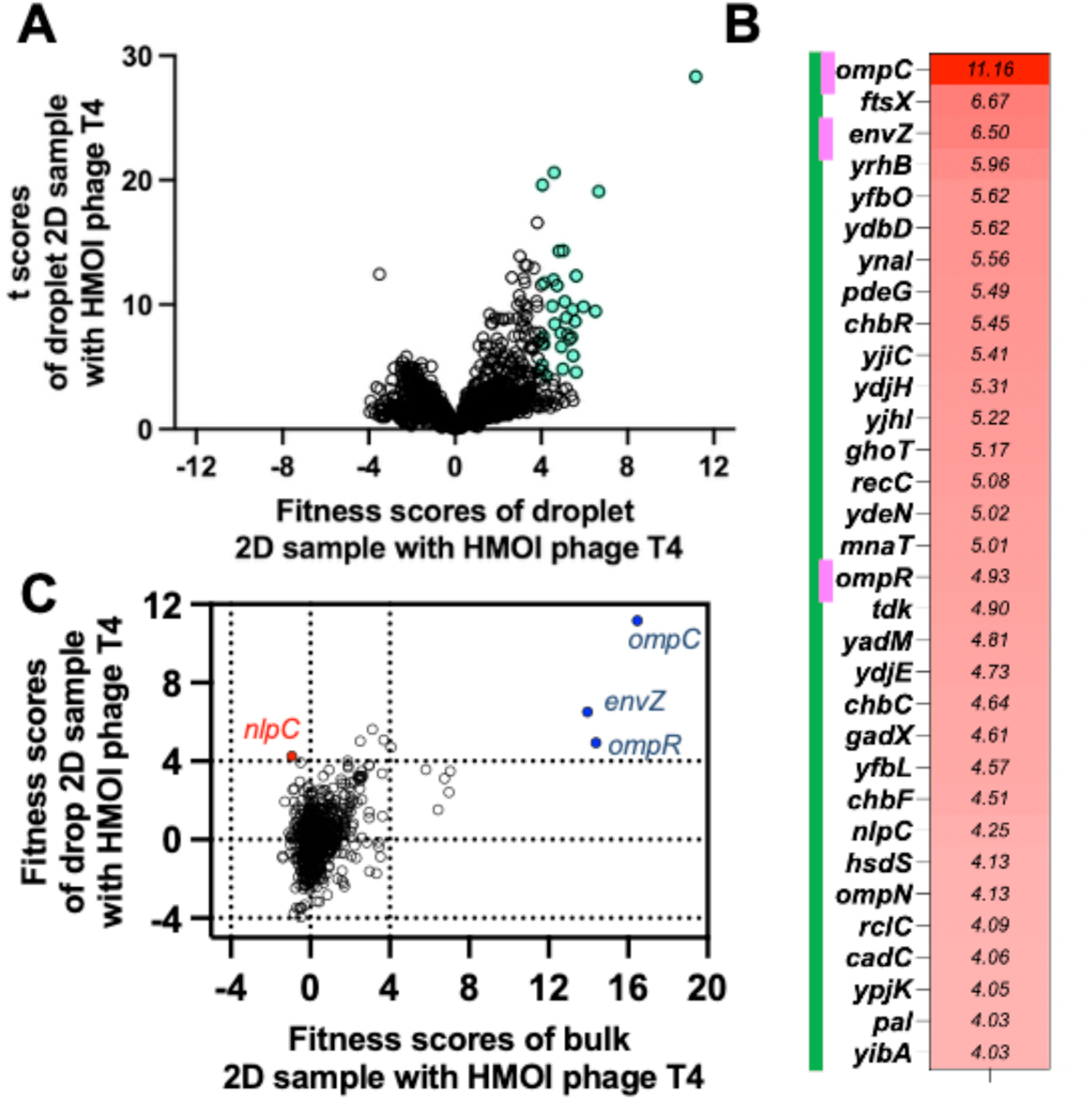
Reveal the resistance of *E. coli* on phage T4 using Droplet RB-TnSeq. (A) Fitness and *t* score of *E. coli* Keio ML9a RB-TnSeq mutants with HMOI phage T4 for 2 days. (B) Genes with |fitness| > 4 and t score > 4. (C) Fitness scores of *E. coli* Keio ML9a mutants exposed to phage T4 from Droplet RB-TnSeq and Bulk RB-TnSeq.

To compare Droplet RB-TnSeq with Bulk RB-TnSeq, we plotted the fitness scores of *E. coli* Keio ML9a mutants exposed to phage T4 from both methods in *Figure 3C*. Only genes with non-zero mutant counts in both methods could be plotted. Most genes from *Figure 3B* are absent here because their abundances in Bulk RB-TnSeq dropped below detection due to microbial competition. From 560,000 reads per sample, only 822 of 3,670 mutants had nonzero counts in Bulk RB-TnSeq, compared with 2,311 in Droplet RB-TnSeq, highlighting the improved resolution of the droplet-based method. Among shared hits, both approaches identified the receptor genes (*ompC*, *envZ*, *ompR*) with fitness >4, and overall results were consistent: mutants with fitness >4 in one method generally showed positive fitness (>1.5) in the other. The exception was *nlpC*, which scored highly in Droplet RB-TnSeq but negatively in Bulk RB-TnSeq (highlighted in red).

To explore the functional roles of genes associated with phage T4 resistance, we mapped those with fitness > 3 and *t* score > 3 to the KEGG BRITE database. A total of 88 genes were assigned to KEGG modules, with 56 linked to *E. coli* metabolisms (*Figure 4A*; full list in *Table S5*). Among these, nine genes belonged to metabolic pathways, followed by five related to two-component systems and four involved in biofilm formation. To gain further insight into resistance mechanisms, we summarized genes related to selected metabolic pathways identified by Droplet RB-TnSeq in *Figure 4B*. Several mutants in the starch/sucrose metabolism pathway, such as *chbR*, *chbC*, and *chbF*, showed high fitness and *t* score during phage T4 infection, suggesting a possible role in resistance. Additionally, genes encoding outer and inner membrane proteins also exhibited high fitness scores, including the known receptor gene *ompC* and other *omp* genes, as well as several novel candidates. Multiple LPS biosynthesis genes also demonstrated moderate resistance (fitness scores between 3 and 4), with higher scores typically observed for genes involved in inner core LPS structures. The contribution of loss of LPS to T4 resistance was validated by the mutants of *rfaC* and *lpcA* (*Figure S4*). Furthermore, two uncharacterized genes, *yfbO* and *yfbL*, showed exceptionally high fitness and *t* scores in Droplet RB-TnSeq.

**Figure 4.**
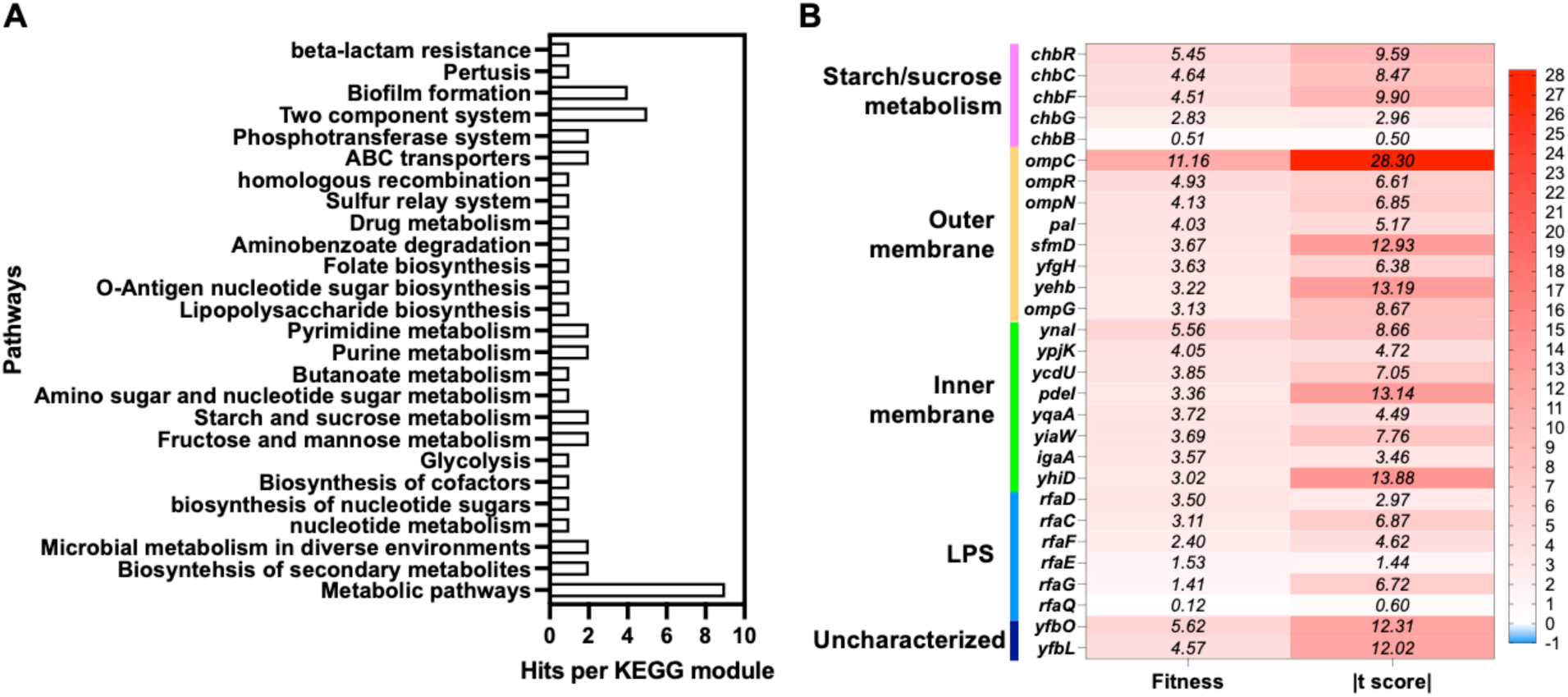
The functional roles of genes associated with phage T4 resistance. (A) A total of 88 genes were assigned to KEGG modules, with 56 linked to *E. coli* metabolisms (B) Genes related to selected metabolic pathways identified by Droplet RB-TnSeq.

### Revealing the resistance of *E. coli* on phage N4 using Droplet RB-TnSeq

Droplet RB-TnSeq was also employed to examine the role of *E. coli* genes in phage N4 infection under the optimized conditions—a high multiplicity of infection (HMOI) and a two-day incubation (2D). *Figure 5A* displays the corresponding fitness and *t* scores of the *E. coli* Keio ML9a RB-TnSeq mutants in the presence of phage N4. Selected genes meeting the criteria of fitness > 4 and *t* score > 4 are marked in green and listed in *Figure 5B*. Consistent with previous findings, known N4 resistance genes such as *nfrA, nfrB, wecB,* and *dgcJ* were successfully identified using Droplet RB-TnSeq (marked pink in *Figure 5B*). In addition, several previously uncharacterized genes emerged as promising candidates potentially involved in phage N4 resistance.

We also compared Droplet RB-TnSeq and Bulk RB-TnSeq in the context of phage N4 infection. *Figure 5C* shows the fitness scores of *E. coli* Keio ML9a mutants with nonzero read counts in both methods. Overall, the results correlated well, indicating strong agreement when mutants were detected by both approaches. However, several mutants identified in Figure 5B, including *yfbL*, *chbC*, and *yibA*, were not detected in Bulk RB-TnSeq. Fitness scores from Droplet RB-TnSeq were generally higher, suggesting that in the absence of competition from highly resistant mutants (*nfrA*, *nfrB*, *wecB*), moderately resistant strains can grow more effectively. For instance, *igaA*, *ygaZ*, *wcaD*, and *dgcQ* mutants exhibited higher fitness in Droplet RB-TnSeq compared to Bulk RB-TnSeq, highlighted in red in *Figure 5C*. A full list of identified genes is provided in *Figure S5*. As expected, both methods captured the known receptor-related genes (*nfrA*, *nfrB*, *wecB*, *dgcJ*) with fitness scores >5. Notably, *dgcQ* mutants exhibited resistance to phage N4 in Droplet RB-TnSeq (fitness = 3.8, *t* score = 3.2) but not in Bulk RB-TnSeq (fitness = –0.3, t score = –0.4). Previous studies have shown that *dgcQ* can function as a backup for *dgcJ* during N4 infection,^26^ underscoring the advantage of Droplet RB-TnSeq in detecting milder resistance phenotypes.

To investigate the biological functions of genes linked to phage N4 resistance, we analyzed those with fitness > 3 and *t* score > 3 using the KEGG BRITE database. This analysis resulted in the assignment of 115 genes to KEGG modules, of which 74 were related to *E. coli* metabolic pathways (*Figure 6A*; complete list in Table S5). This number is substantially higher than that identified for phage T4, suggesting that a broader range of cellular mechanisms may contribute to resisting or slowing phage N4 infection. This observation is consistent with previous reports indicating that the lytic phage T4 has a shorter latent period than phage N4,^27–29^ making phage T4 more aggressive and rapidly lytic in comparison with phage N4. In *Figure 6A*, sixteen genes were categorized under metabolic pathways, followed by five linked to ABC transporters, five to biofilm formation, and four to lipopolysaccharide (LPS) biosynthesis.

**Figure 5.**
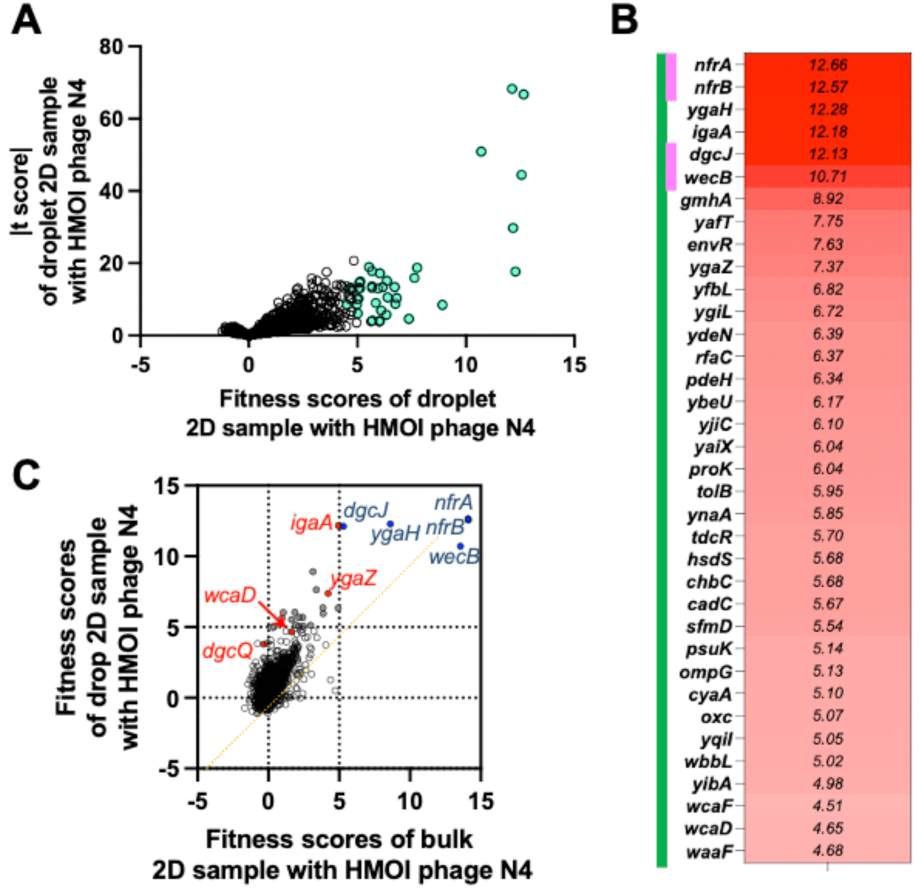
Revealing the resistance of *E. coli* on phage N4 using Droplet RB-TnSeq. (A) Fitness and *t* score of *E. coli* Keio ML9a RB-TnSeq mutants with HMOI phage N4 for 2 days. (B) Genes with |fitness| > 4 and *t* score > 4. (C) Fitness scores of *E. coli* Keio ML9a mutants exposed to phage N4 from Droplet RB-TnSeq and Bulk RB-TnSeq.

**Figure 6.**
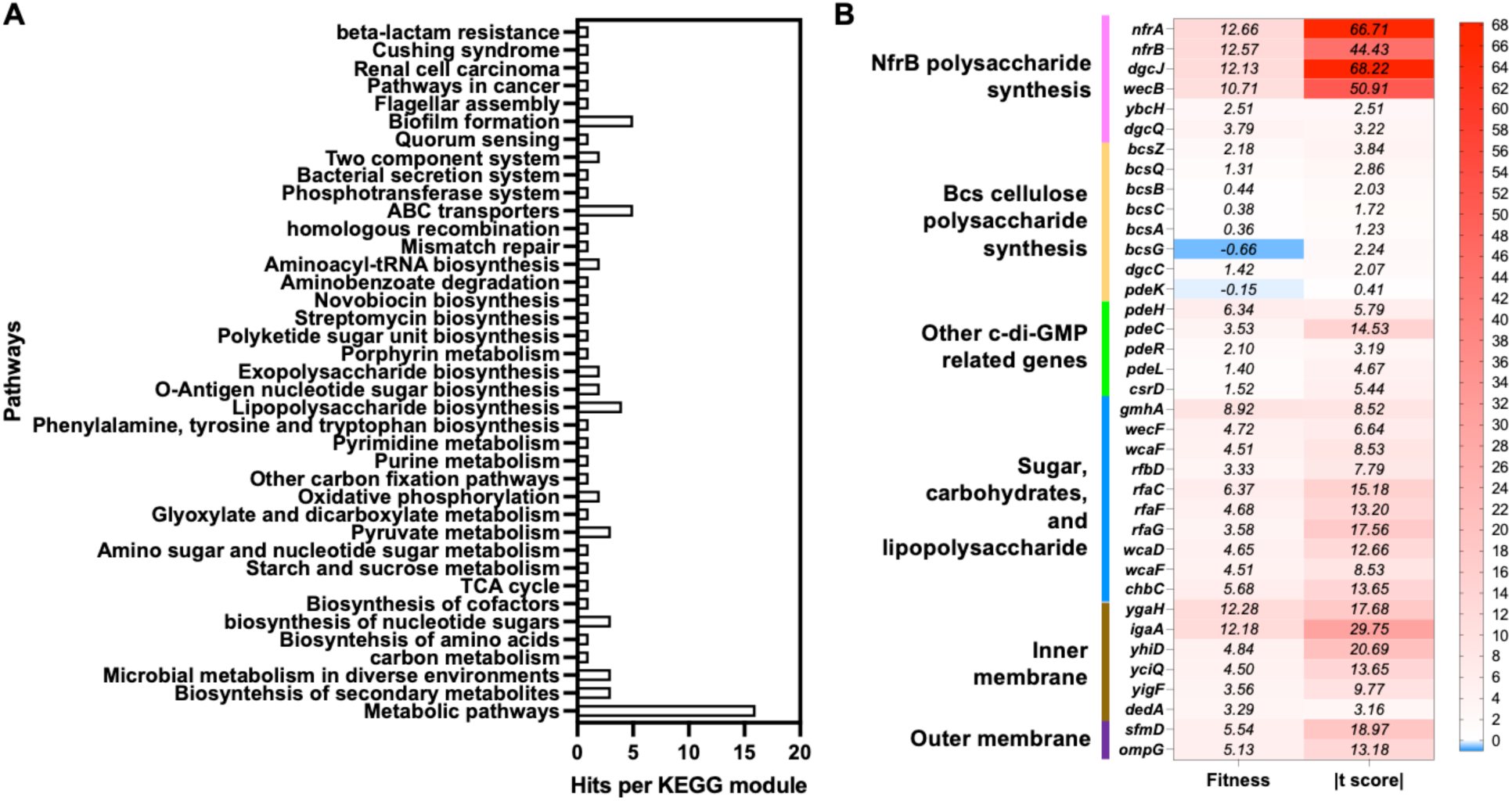
The functional roles of genes associated with phage N4 resistance. (A) A total of 115 genes were assigned to KEGG modules, with 74 linked to E. coli metabolisms (B) The fitness and *t* scores of genes involved in selected metabolic pathways.

To gain further insights into resistance mechanisms, we summarized the fitness and *t* scores of genes involved in selected metabolic pathways—some of which are potentially related or known not to be involved in N4 resistance—in *Figure 6B*. The Droplet RB-TnSeq results confirm the NfrB polysaccharide synthesis pathway as the entry route for phage N4, which is reported in previous studies^26,30^ highlighting the essential roles (high fitness score) of *nfrA*, *nfrB*, *wecB*, and *dgcJ*, along with the partial contributions (moderate fitness score) of *ybcH* and the minor, non-essential role (moderate fitness score) of *dgcQ* in the pathway. Additionally, our Droplet RB-TnSeq analysis shows that the Bcs cellulose polysaccharide synthesis pathway—despite its similarity to the NfrB pathway—is not involved in phage N4 infection, consistent with prior findings.^31,32^ C-di-GMP signaling is another pathway associated with phage N4 resistance. Droplet RB-TnSeq identified several related genes, such as *dgcJ* and *pdeH*, as contributing to resistance. Mutants of *dgcQ* and *pdeC* also showed moderate resistance, while *pdeL* and *csrD* mutants did not display significant fitness increases during phage N4 infection. Several genes involved in sugar, carbohydrate, and lipopolysaccharide biosynthesis pathways showed notably high fitness scores, suggesting a key role in resistance. These include *wecF* (related to enterobacterial common antigen), various *wca* genes (involved in colanic acid synthesis), and LPS biosynthesis genes such as *gmhA*, *rfbD*, and members of the *rfa* cluster. Finally, several genes encoding inner and outer membrane proteins, including *ygaH*, *igaA*, *sfmD*, and *opmG*, also showed elevated fitness scores, suggesting their possible involvement in phage N4 infection.

### Validation of candidate genes associated with phage T4 and N4 infection identified by Droplet RB-TnSeq

To validate the phage resistance phenotypes identified by Droplet RB-TnSeq, we conducted phage plaque assays using individual gene deletion mutants from the *E. coli* K-12 BW25113 Keio collection. The *E. coli* K-12 BW25113 wild-type strain was included as a control. Based on their significance in both T4 and N4 resistance screens and the availability in the Keio library, we selected 11 mutants (*ygaZ*, *ygaH*, *chbR*, *chbC*, *chbF*, *ycjQ*, *waaD*, *waaF*, *yibA*, *yfbO*, *yfbL*) for testing against phage N4, and 9 mutants (*ompC*, *narW*, *waaC*, *nlpC*, *chbF*, *chbC*, *chbR*, *letA*, *waaF*) for testing against phage T4. The *ompC* mutant served as a negative control for phage T4. All selected Keio mutants were confirmed by PCR. Phage resistance was evaluated using a standard plaque assay. Briefly, bacterial lawns were prepared on soft agar plates, followed by serial dilutions of either phage N4 or T4 applied directly onto the surface. Plaques (zones of clearance) indicated successful phage infection. Results are shown in *Figure 7*. Nearly all mutants tested for phage N4 resistance displayed clear resistance phenotypes, with many exhibiting more than a two-log reduction in plaque formation. In contrast, the mutants tested for phage T4 resistance only showed a marginal difference from the wild-type strain. The possible reason is discussed in the discussion. As a result, we focused our follow-up validation on N4-resistant candidates.

**Figure 7.**
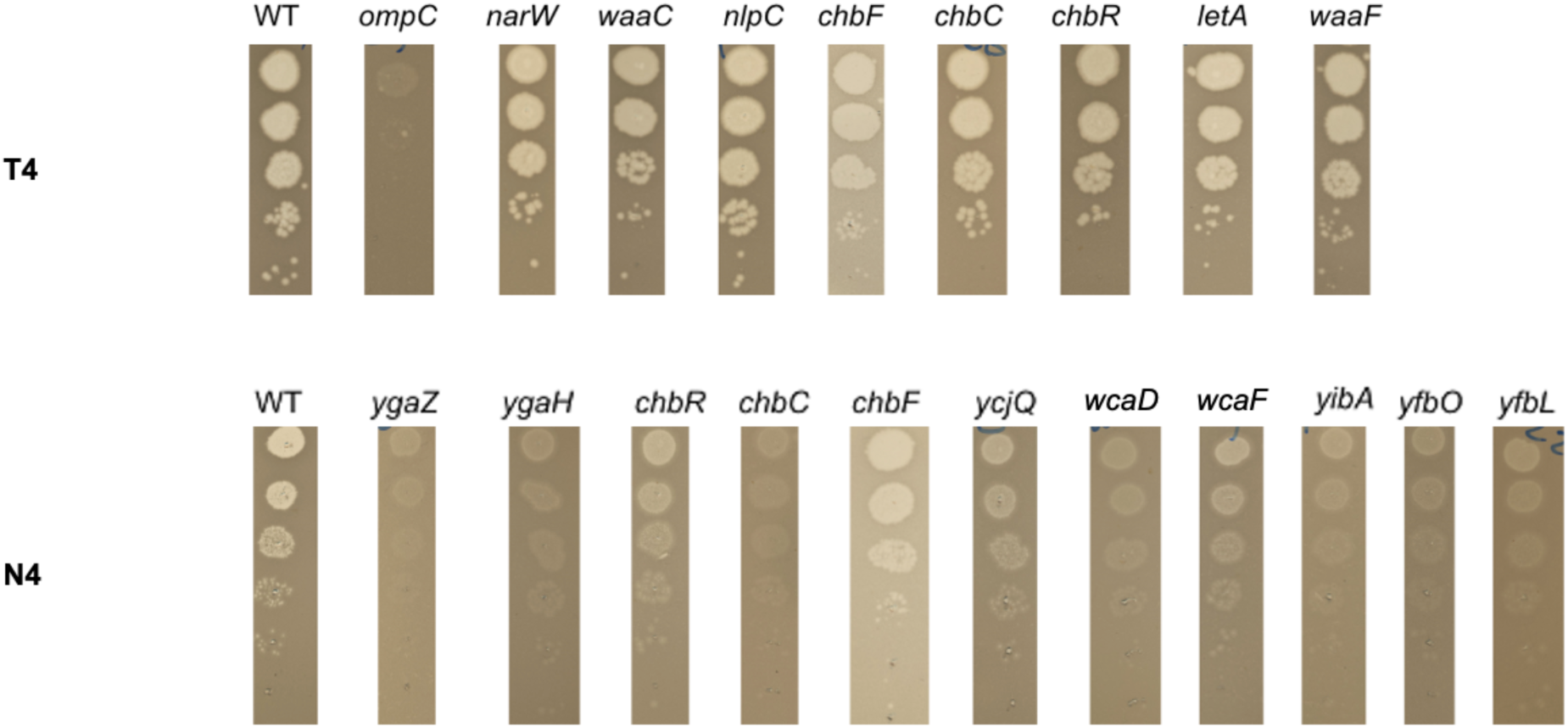
Experimental validation of top-scoring gene hits for phage T4 and N4 resistance found in Droplet RB-TnSeq using efficiency of plating experiments. WT is the wild type *E. coli* Keio library strain BW25113. And the mutants are picked from the *E. coli* Keio mutant library.

For further investigation, we selected *chbR*, *chbC*, *chbF*, *ygaZ*, *ygaH*, *yfbO*, and *yfbL*, which clustered together and showed strong phage N4 resistance in plaque assays. To confirm their role in resistance, we performed genetic complementation using plasmids from the ASKA library. Each mutant was transformed with its corresponding gene on the pCA24N vector via electroporation. Complementation was confirmed by PCR and gel electrophoresis. The plasmid and primers are shown in *Table S3* and *S4*. As controls, each mutant and the wild-type strain were also transformed with an empty plasmid. Results are shown in *Figure 8*. Three experimental groups were included: (1) wild-type strain with empty plasmid (WT), (2) mutants with empty plasmid (-), and (3) mutants with complemented plasmid (+). All strains were tested for phage N4 susceptibility using plaque assays. As expected, the WT strain with empty plasmid remained susceptible to phage N4, while mutants with empty plasmids exhibited resistance. Upon complementation, all mutants restored susceptibility to phage N4, confirming that disruption of *chbF*, *chbC*, *chbR*, *yfbL*, *yfbO*, *ygaZ*, and *ygaH* directly contributed to the resistance phenotype.

**Figure 8.**
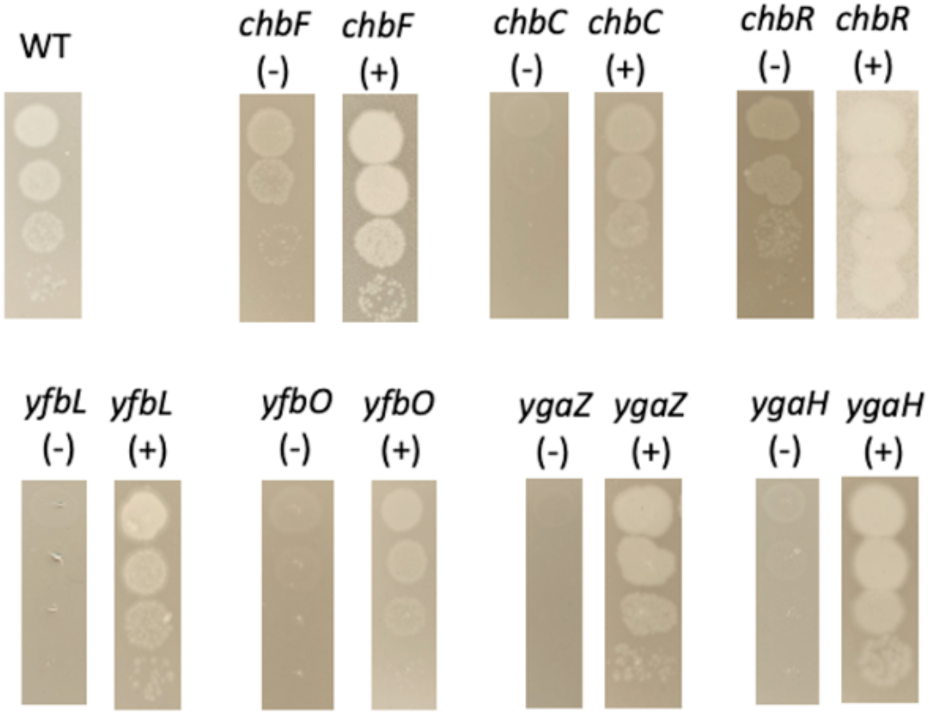
Experimental validation of top-scoring gene hits for phage N4 resistance found in Droplet RB-TnSeq using efficiency of plating experiments. WT is the wild type *E. coli* Keio library strain BW25113. We used ASKA plasmid as the complementation vector. (-) represents the mutant strain with control vector pCA24N. (+) represents the mutants with the complement of the mutant gene.

To assess whether these mutations confer cross-resistance to other phages, we conducted a broader screen against phages T3, T5, T6, T7, P1, Lambda, and LZ4. The same set of validated phage N4-resistant mutants (*chbF*, *chbR*, *chbC*, *ygaZ*, *ygaH*, *yfbL*, *yfbO*) from the *E. coli* K-12 BW25113 Keio collection and the wild-type strain *E. coli* K-12 BW25113 were subjected to plaque assays. However, none of the mutants exhibited significant resistance to these additional phages (data not shown), indicating that their resistance is specific to phage N4.

## Discussion

Three novel gene clusters associated with phage N4 resistance were identified through Droplet RB-TnSeq and subsequently validated via phage plaque assays and gene complementation: the *chb*, *yga*, and *yfb* gene clusters. The *E. coli chb* operon is known to mediate the transport and metabolism of chitobiose and related N-acetylglucosamine (GlcNAc)-containing oligosaccharides.^33^ Specifically, ChbC is a component of the inner membrane permease complex that imports chitobiose and phosphorylates it, resulting in intracellular accumulation of chitobiose-6-phosphate. *chbF* encodes a phospho-β-glucosidase that hydrolyzes chitobiose into GlcNAc-phosphate units, while ChbR functions as a dual-purpose AraC family transcriptional regulator that modulates the operon in response to sugar availability. We did not detect any barcode reads for the mutants of other *chb* operon genes such as *chbA*, *chbB*, and *chbG*, likely due to limitations in library coverage and sequencing depth. For instance, *chbG* yielded zero reads even in the time-zero sample, and the remaining *chb* genes showed fewer than 30 reads in the time-zero sample and zero reads in phage-treated conditions. Future experiments with larger volumes of droplet cultures could improve the coverage issue. Although the precise mechanism by which *chb* genes influence phage N4 infection remains unclear, we hypothesize that their disruption may alter cell surface sugar composition or outer membrane integrity, thereby affecting phage recognition or entry. The *ygaZ* and *ygaH* genes form a putative two-component transporter system localized in the inner membrane and have been implicated in L-valine export. Overexpression of this system has been shown to enhance L-valine production,^34^ while deletion of *ygaH* has been associated with resistance to the antimicrobial agent ornidazole.^35^ Despite these insights, functional data on *ygaZ* and *ygaH* remain limited. Even less is known about the *yfbL* and *yfbO* genes. *yfbL* is predicted to encode a peptidase,^36^ whereas YfbO is annotated as an uncharacterized protein with no defined function. Further investigation into their roles in phage N4 infection may provide novel insights into previously unknown host factors involved in bacteriophage resistance.

The phage plaque assay did not fully validate the top hits identified from phage T4 infection by Droplet RB-TnSeq (*Figure 7*). This discrepancy likely arises from two factors. First, Droplet RB-TnSeq is a sequencing-based method that tracks the presence of genetic markers rather than cell viability. Even if cells are ultimately killed by phage T4, the abundance measured reflects the maximum population size reached prior to lysis. In such cases, increased fitness scores do not necessarily indicate true resistance, but rather faster growth or delayed killing. Although this still provides valuable insight into bacterial defense mechanisms, it may not align with outcomes from phage plaque assays, which directly assess susceptibility versus resistance in a binary manner. Given that phage T4 is more aggressive and rapidly lytic than phage N4, it is likely that many host responses only delay cell death rather than prevent infection. In this case, moderate resistance corresponds to a delay in killing at the tested MOI—this feature would not be captured by plaque assays. Second, Droplet RB-TnSeq maintains a relatively constant selective pressure, as droplets separate each individual cell which prevents an increase in MOI during infection. By contrast, in plaque assays, cells lysed early release additional phage, amplifying MOI. This escalating pressure eliminates moderately resistant cells that would otherwise persist at the original MOI. Since T4 lyses *E. coli* within minutes, this effect is especially pronounced. Especially for phage plaque asay, when phage spots are still wet on agar plates, rapid diffusion of the new released phages further accelerates spread, intensifying selective pressure. Consequently, plaque assays are poorly suited to validate moderately/marginally resistant mutants, particularly against highly lytic phages such as T4.

Droplet RB-TnSeq reveals the genes that have non-essential roles during phage infection. For example, *dgcQ* shows strong fitness advantages in Droplet RB-TnSeq with N4 (fitness and t-score > 3; Figure 6B), yet was not identified in Bulk RB-TnSeq (fitness = –0.3). Junkermeier et al.^26^ demonstrated that *dgcQ* can function as a backup for *dgcJ* during N4 infection, which agree with our Droplet RB-TnSeq results. Similarly, the plaque assay of *dgcQ* mutant revealed only a marginal defect under phage N4 infection compared to wild type.^26^

Droplet RB-TnSeq, which integrates droplet microfluidics with barcoded transposon sequencing, represents one of the highest-throughput experimental platforms currently available for investigating bacteria–phage interactions and associated gene functions. This method enables the parallel execution of millions of individual “reactions” within droplets and allows for the simultaneous readout of all outcomes through a single sequencing run. In addition to revealing and validating the role of *chb*, *yga*, and *yfb* gene clusters on phage N4 infection, this approach revealed numerous single-gene mutants with high fitness scores during phage T4 or N4 infection. Further investigation of these candidate genes may uncover new mechanisms underlying bacterial responses to phage infection. Beyond host–phage interaction studies, Droplet RB-TnSeq can also be adapted to explore a range of other phenotypes, including antibiotic resistance and nutrient utilization, by simply replacing the phage input with antibiotics or specific nutrients. We are actively developing such applications, which promise to become powerful tools for generating large-scale experimental data.

Despite its strengths, Droplet RB-TnSeq also has several limitations in studying bacteria–phage interactions. First, the method relies on a transposon single-gene mutant library, which restricts the types of genetic effects that can be observed to the host factors. Expanding the scope to include gain-of-gene approaches—such as Dub-seq^37^ or Boba-seq^38^—or double-mutant libraries would provide a more comprehensive view of host–phage interactions. Second, this approach infers phage susceptibility by measuring cell death or growth defects, making it suitable primarily for lytic phages, but not for lysogenic ones that do not immediately kill the host. Third, as mentioned above, RB-TnSeq estimates mutant fitness based on the abundance of their genetic barcodes, which reflects overall cell count without distinguishing between live and dead cells. While generally reliable, this measure may be confounded by dynamic host–phage interactions. For example, some mutants may delay lysis without ultimately surviving the infection.

## Conclusion

Systematically annotating gene functions and rapidly identifying bacterial host factors involved in phage infection remains a significant challenge. Traditional approaches such as Random Barcode Transposon-Site Sequencing (RB-TnSeq) and Transposon-Site Sequencing (Tn-Seq), while powerful, often introduce strong microbial interactions and selection pressures that can obscure host factors beyond surface receptors. To overcome these limitations, we employed a droplet-based RB-TnSeq method, which encapsulates individual bacterial strains within microdroplets. This approach retains the key advantages of RB-TnSeq—such as high throughput and cost-effectiveness—while minimizing microbial competition and MOI escalation during phage challenge. We optimized the Droplet RB-TnSeq protocol by tuning droplet size, cell loading density, and culture duration, and demonstrated that the method is both reliable and reproducible. Using this approach, we successfully recovered known resistance mechanisms against phages T4 and N4 and additionally identified and validated several novel genes potentially associated with resistance to both phages. Focusing further on N4, we confirmed that loss of the following genes - *chbR*, *chbC*, *chbF*, *ygaZ*, *ygaH*, *yfbL*, or *yfbO* - confer phage N4 resistance. Overall, Droplet RB-TnSeq provides a powerful and complementary tool for identifying bacterial host factors beyond classical phage receptors, offering new insights into bacteria–phage interactions and expanding the scope of functional genomics in microbial systems.

## Methods

### Materials, strains, and growth conditions

Bacterial strains, bacteriophages, plasmids, and primers used in this study are listed in Table S1, S2, S3, and S4, respectively. All the oligonucleotides were purchased from Integrated DNA Technologies (IDT). Plasmids and primers were stored at -20°C. Bacterial strains were grown in Luria Bertani (LB) Lennox medium containing 10 g/l tryptone, 5 g/l yeast extract, and 5 g/l sodium chloride (NaCl) with or without appropriate antibiotics at 37°C with 200 rpm shaking. The bacterial strains were stored at -80°C in 15% glycerol stock. The *E. coli* Keio ML9a RB-TnSeq library strains (Generation 0, 1 mL) were obtained from Deutschbauer Lab at Lawrence Berkeley National Laboratory (LBNL) and cultivated at 37°C with 200 rpm shaking for overnight by mixing with 100 ml LB Lennox medium supplemented with 25 ug/ml Kanamycin in a flask. The overnight culture was aliquoted into 1.5 mL stocks to make 15% glycerol stocks (Generation 1) and stored at -80°C until use.

All the strains were re-inoculated into the appropriate medium from the glycerol stocks and cultured at 37°C with 200 rpm shaking overnight before use. *E. coli* K-12 BW25113 was cultured in LB Lennox medium; *E. coli* deletion mutations from Keio mutation library were cultured in LB Lennox medium supplemented with 50 ug/ml Kanamycin; *E. coli* ASKA library strains were cultured in LB Lennox medium supplemented with 50 ug/ml Chloramphenicol and 0.1 mM Isopropyl beta-D-1-thogalactopyranoside (IPTG); and *E. coli* complemented strains were constructed in this study and cultured in LB Lennox supplemented with 50 ug/ml Kanamycin, 50 ug/ml Chloramphenicol and 0.1 mM IPTG. The *E. coli* Keio ML9a RB-TnSeq library strains were cultured from ML9a glycerol stock (1.5 ml, generation 1) in LB Lennox supplemented with 50 ug/ml Kanamycin at 37°C with 200 rpm till the OD_600_ reached 1 (exponential phase). Then the ML9a RB-TnSeq library strains were diluted to desired concentration for the droplet-based phage infection experiments.

All the bacteriophages were propagated on *E. coli* BW25113 strain. Standard protocols were followed for the phage propagation. The propagated bacteriophages were routinely sterilized by 0.22 um filter and stored in a modified SM buffer [SM buffer (Teknova) supplemented with 10 mM Calcium Chloride and Magnesium Sulphate (Sigma)] at 4°C. After propagation and before each experiment, the phage titer was measured by the bacteriophage plaque assay. Briefly, host strain soft agar plates were first made by pouring the mixture of 100 ul of the overnight culture of host strain *E. coli* BW25113 with 4 ml warm (56°C) 0.7% LB agar into a petri dish. After the host strain soft agar plates cooled down and solidified, 5 ul of phage stock was spotted on the host strain soft agar plates with 3 replicates by 10-fold serial dilution. The phage titer was calculated based on the phage plaque after incubation at 37°C.

### Microfluidic devices

The microfluidic devices were fabricated by four steps – (1) designing a photo mask, (2) making a master mold, (3) fabricating a polydimethylsiloxane (PDMS) microfluidic device, and (4) performing surface treatment of microfluidic channels. First, the microfluidic device photo mask was designed by AutoCAD (AutoDesk Inc.). The design of our microfluidics device is shown in Figure S1 and the dwg file is available in Supplementary files. The photo mask was fabricated by FineLine Imaging Inc. Second, the master mold was made by a negative photoresist (SU-8 3025, MicroChem) on a 4” silicon wafer (Prime grade, WaferNet) followed by the SU-8 photolithography using the designed photo mask. The protocol was provided by MicroChem Inc. Third, the PDMS microfluidic devices are fabricated by pouring Sylgard 184 silicone elastomer kit (1:10 ratio of base: curing reagent, degassed, Dow Corning Inc.) on the master mold. Then the PDMS microfluidic device was cured at 60°C overnight. After punching the inlet and outlet, the PDMS microfluidic device and a glass slide were bonded together to assemble to a microfluidic chip followed by the oxygen plasma treatment (Jelight UV Ozone Surface Modification System). In addition, the microfluidic chips were treated by Aquapel (PPG industries) and baked at 65°C for 10 min to make the channel surface hydrophobic before they are ready to use.

### Droplet-based RB-TnSeq assay

Single cell bacterial strain was mixed with bacteriophages in the microfluidic chip (Figure 1). Two aqueous suspensions – bacterial RB-TnSeq library strains and bacteriophages – were loaded into 3 ml plastic syringes (BD), respectively. The syringe containing bacterial strains was equipped with a disc stirrer (5mm, V&P Scientific VP 772DP-N42-5-2) that was powered by the magnetic stirring system (V&P Scientific VP 710D3) during the droplet generation to make the bacterial suspensions homogeneous. Oil HFE-7500 with 2%wt 008-FluoroSurfactant (RAN biotechnology) was loaded into a 10 ml plastic syringe (BD). The bacterial RB-TnSeq library strain – *E. coli* ML9a was cultured in LB Lennox medium supplemented with 50 ug/ml Kanamycin to OD_600_ = 1, and diluted by 2X LB Lennox medium to OD_600_ = 0.001 as the input of the bacterial RB-TnSeq library strains during droplet culture generation. 1 ml of the input was taken as the Time0 sample. After centrifuge at 10000 g for 5 min, stored immediately at -80°C until DNA extraction. Three phage concentrations were used in the droplet culture generation: no phage (No-phage sample), 4.4×10^7^ pfu/ml (low-phage sample) and 5.1×10^9^ pfu/ml (high-phage sample). The phage suspension was diluted from ∼ 5 x 10^11^ pfu/ml phage stock in SM buffer with 10 mM CaCl_2_ and _MgSO4_. The phage stock was stored at 4°C and the phage concentration was confirmed by plaque assay before use.

Droplet cultures with ∼ 60 um diameter were generated using the microfluidic chip. Each droplet culture contained the bacterial mutation strain and phages. There was 1 ml of the input bacterial suspension (∼ 1.77 x 10^6^ cells/ml, OD_600_ ≈ 0.001) used in each condition. The droplet cultures were incubated in a 50 ml falcon tube at 37°C with 80 rpm shaking to allow the microbial cell to grow with phages for 1 day (1-day samples) and 2 days (2-day samples).

The number of *E. coli* strains and phages in each droplet is determined by the concentration of *E. coli* Keio ML9a RB-TnSeq library, the concentration of *E. coli* phage, and the droplet size. It is described by Poisson distribution.^39^

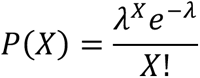

Where, *P(X)* is the percentage of the droplets that contain X cells. λ is the overall average number of cells per droplet that can be calculated by multiplying the input cell concentration by the droplet volume. The droplet volume is calculated by the droplet diameter since the droplets are perfect spheres. The droplet sizes were controlled by the dimension of the microfluidic chip and flow rates of samples and oil. In our experiment, we achieved a single microbial cell per droplet with the average ratio of phage to bacteria (MOI) that is either 5 (phage density 4.4×10*^7^* pfu/ml) or 580 (phage density 5.1×10^9^ pfu/ml). The simulated distribution of *E. coli* cells and phage cells per droplets are shown in Figure S2.

After incubation, the droplet samples were de-emulsified by removing the bottom oil and adding 100 ul *1H, 1H, 2H, 2H-*perfluoro-1-octanol (Sigma). After gently vortexing, the water layer was separated from the oil layer. The water layer containing bacterial cells was collected. Cells were pelleted by centrifuge at 10000 g for 3 min at 4°C, supernatant was carefully removed, and the pellet was stored at -80°C until DNA extraction. The whole process was performed on ice and completed in 8 min to minimize the phage lysis during the sample process. For each bacterial RB-TnSeq library strains and phage, there were three samples (no-phage, low-phage, and high-phage), three time points (Time 0, 1-day, and 2-day), and two replicates. The cell pellets were stored at - 80°C.

### Barseq and data analysis

We extracted genomic DNA (gDNA) of the cell pellet samples using Qiagen Blood and Tissue DNA extraction kit. We used the BarSeq PCR protocol described before^1^ using 50 ul PCR reaction with 20 umol of each BarSeq PCR primer (Table S4) and 150 ng of template gDNA. For HiSeq4000 sequencing, we used an equimolar mixture of BarSeq_P2 primers and new BarSeq3_P1 primers. The BarSeq_P2 primer carries the tag required for demultiplexing by Illumina software, while the BarSeq3_P1 primer includes an additional sequence to verify that reads originated from the correct sample. Specifically, the BarSeq3_P1 primer is identical to the previously described BarSeq_P1 sequence, with 1–4 variable N’s (depending on the index), followed by the reverse (not the reverse complement) of the six-nucleotide index sequence. This modification was introduced to eliminate barcode bleed-through and to improve cluster and sample discrimination on the HiSeq4000 platform. All experiments performed on the same day and sequenced on the same lane were treated as a “set.” Equal volumes (5 μl) of individual BarSeq PCR products were pooled, and 50 μl of the pooled product was purified using the DNA Clean and Concentrator kit (Zymo Research). Libraries were eluted in 40 μl of water and sequenced on an Illumina HiSeq4000 instrument with 50-cycle single-end runs. Typically, 96 BarSeq samples were multiplexed per lane for both RB-TnSeq and Dub-seq libraries.

BarSeq fitness data were analyzed as previously described.^15^ The method is documented by Morgan Price (LBNL) in https://genomics.lbl.gov/supplemental/strongselection/Keio/. Briefly, the fitness of each mutation strain is log2(strain barcode abundance after incubation/strain barcode abundance at Time0). The fitness of each gene is the weighted average of the fitness of its mutation strains. A t-like test statistic (t-score) is used for identifying the confidence. The *t*-score is defined as the gene fitness divided by the standard error. And the standard error is the maximum of two estimates: the consistency of the fitness for the mutation strain in that gene and the number of reads for the gene. High absolute *t*-score indicates the high confidence of the fitness results. In this study, we remove all the strains with zero reads in the results.

### Experimental validation of individual phage resistance phenotypes

A selective set of high phenotype genes were validated by the phage plaque assay. First, we used single gene knockout strains to validate the phenotype we obtained from Droplet RB-TnSeq assay. Single gene knockout strains were taken from the KEIO collection^40^ and the mutations were verified via PCR for correct insertion of the KanR gene using primers oAH1-oAH29 (*Table S3*). Each single gene knockout strain was cultured at LB medium supplemented with 50 ug/ml overnight. Then the plaque assay was performed as shown above. Briefly, bacterial soft agar plates were first made by pouring the mixture of 100 ul of the overnight culture of the single gene knockout strain with 4 ml warm (56°C) 0.7% soft LB agar into a petri dish. After the host strain soft agar plates cooled down and solidified, 5 ul of phage stock was spotted on the host strain soft agar plates with 3 replicates by 10-fold serial dilution. The phage titer was calculated based on the phage plaque before use. The soft agar plates with bacteria and phages were incubated at 37°C until the plaques were clearly visible. The phage resistance of the single gene knockout strain was calculated by the plaque shown on the plates. More phage plaque indicates the high susceptibility to phage.

To confirm the phage resistance phenotype is caused by the gene that was knocked out, we constructed the gene complemented strains to investigate if the phenotype could be restored after the gene complementation. First, single gene knockout strains were taken from the KEIO collection and grown in Miller LB with Kanamycin (50 ug/mL) for 12-18 hours, followed by a 1:1000 dilution and subculture to mid-exponential phase (2-3 hours). The cells were made electrocompetent by four wash cycles in chilled 10% glycerol. Each KEIO strain was verified via PCR for correct insertion of the KanR gene using primers oAH1-oAH29 (*Table S4*). Second, overexpression strains for each gene were then taken from the ASKA collection, which contain variants of the IPTG-inducible, GFP-negative pCA24N plasmid.^41^ The selected strains were grown for 12-18 hours in LB with Chloramphenicol (30 ug/mL) and miniprepped using the NEB Monarch miniprep kit. These ASKA plasmids were sequence verified using sequencing primers oAH30-oAH31 (*Table S3*). Third, the plasmids were transformed via electroporation into the KEIO strains, along with an empty pCA24N vector, and selected using LB with Kanamycin (50 ug/mL) and Chloramphenicol (30 ug/mL). To confirm the gene function, we performed the phage plaque assay on host strain (*E. coli* BW25113) with empty plasmid, single gene knockout strain (KEIO mutants) with empty plasmid, and complemented strains (constructed in this study) with complement genes. Less phage plaque indicates more resistance to phage.

## Acknowledgements

We thank Adam M. Deutschbauer (LBNL) for providing the *E. coli* RB-TnSeq library and Morgan Price (LBNL) for helping with the analysis of RB-TnSeq data. This work conducted by ENIGMA-Ecosystems and Networks Integrated with Genes and Molecular Assemblies (https://enigma.lbl.gov/), a Scientific Focus Area Program at Lawrence Berkeley National Laboratory, was supported by the Office of Science, Office of Biological and Environmental Research, of the U. S. Department of Energy under Contract No. DE-AC02-05CH1123.

## Author contributions

FS, VKM, and APA conceived the project. FS, AH, MM, and VKM conducted the experiments. APA supervised the project. FS analyzed the data. FS and AH wrote the original draft. FS, AH, VKM, and APA reviewed and revised the manuscript.

## Competing interests

APA is a shareholder in and advisor to Nutcracker Therapeutics.

## Data availability

The top hits from the screening are listed in Table S5. The raw Barseq results are available from https://morgannprice.org/FEBA/Keio/.

---

**Figure S1.**
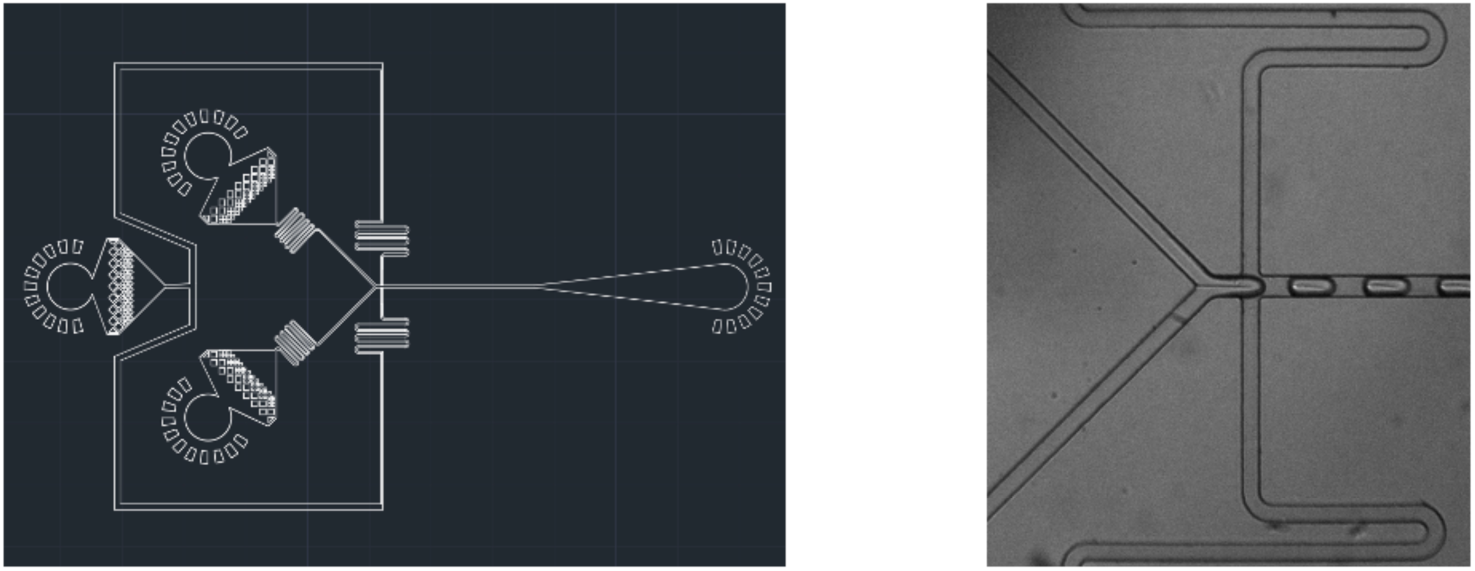
(A) Design of the two inlets mixture droplet generation chip. The chip is fabricated using PDMS on the patterned silicon wafer SU-8 made by SU-8 photolithography. (B) Droplet generation with mixing of bacteria and phages in the chip.

**Figure S2.**
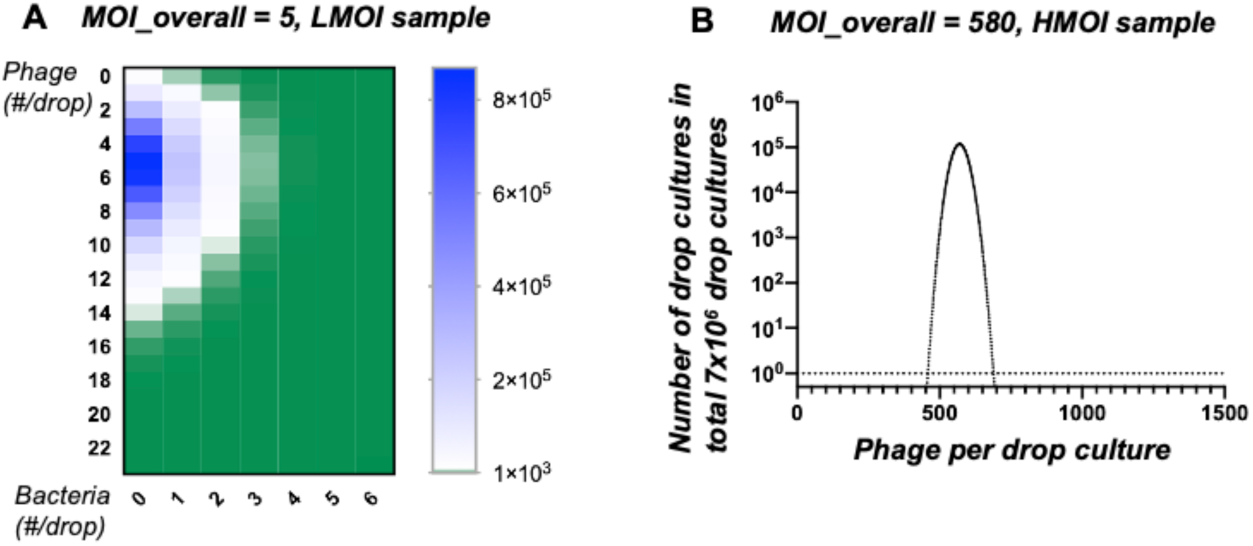
The distribution of phage and bacteria number across droplets used in this study calculated by the Poisson process. (A) The distribution of droplet numbers when the average MOI of phage and bacteria input into the droplet generation is equal to 5. The heatmap displays the numbers of droplets with certain numbers of bacterial cells and phages in total of 7 x 10^6^ generated droplets. The experiment is achieved by encapsulating 1.77×10^6^ cfu/mL bacterial cells and 4.4×10^7^ pfu/mL phage into 60 um diameter droplets. Blue indicates a high number of droplets with a given bacterial/phage combination, while dark green indicates zero droplets. (B) The MOI distribution of droplets when the average MOI is equal to 580. The plot shows the theoretical distribution of phage numbers with single bacterial cells in droplets. The experiment is achieved by encapsulating 1.77×10^6^ cfu/mL bacterial cells and 5.1×10^9^ pfu/mL phage into 60 um diameter droplets.

**Figure S3.**
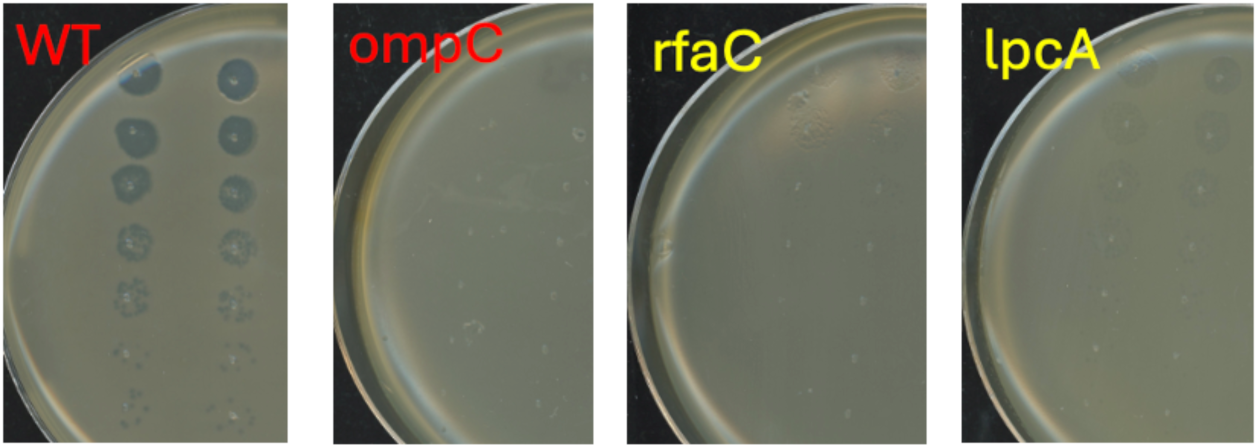
Experimental validation of LPS related genes for phage T4 resistance using efficiency of plating experiments. WT is the wild-type *E. coli* Keio library strain BW25113. And the mutants are picked from the *E. coli* Keio mutant library.

